# Quantitative detection of plant signaling peptides utilizing ELISA

**DOI:** 10.1101/2024.06.27.600388

**Authors:** Maurice König, Zarah Sorger, Gunther Doehlemann, Johana C. Misas Villamil

**Affiliations:** Institute for Plant Sciences, University of Cologne, Cologne, Germany; Cluster of Excellence on Plant Sciences (CEPLAS), University of Cologne, Cologne, Germany

**Keywords:** ELISA, Zip1, peptide detection, phytocytokines, plant signaling peptides

## Abstract

Plant signaling peptides, also known as phytocytokines, are involved in a number of signaling mechanisms, including cell-to-cell communication during plant development and immunity. The detection of small peptides in plant tissues is challenging and often relies on time-consuming and cost-intensive approaches. Here, we present an ELISA-based assay as a rapid and cost-effective method for the detection of naturally released peptides in plant tissues. Our ELISA-based method was developed to detect Zip1, a 17-amino-acid phytocytokine derived from *Zea mays* that elicits salicylic acid signaling in maize leaves. Using a custom peptide-antibody, we designed an experimental pipeline to achieve peptide specificity, selectivity and sensitivity allowing the detection of the Zip1 peptide in complex biological samples. As a proof of concept, we transfected maize protoplasts to overexpress the precursor molecule PROZIP1 and treated maize leaves with salicylic acid to induce native PROZIP1 expression and Zip1 release. Using ELISA, we were able to quantify native Zip1 signals with a detection limit in the nanogram range, which allowed us to detect different Zip1-containing peptides in plant material. This method can be adapted for the detection and quantification of a variety of plant signaling peptides.

## Introduction

Plant endogenous signaling peptides, known as phytocytokines, play an important role in plant growth, development and stress responses by fine-tuning hormonal responses (Gust *et al*. 2017, Hou *et al*., 2021, Rzemieniewski *et al*, 2022). One prominent example is Zip1 (*Zea mays* immune peptide 1), a maize-specific, 17 amino acid peptide that enhances salicylic acid-associated defense signaling upon the release from its precursor PROZIP1 via proteolytic processing (Ziemann *et al*., 2018). The expression of phytocytokine precursors is tightly timed and precisely regulated (Huffaker *et al*., 2006, 2011) and it has been shown to be commonly induced by the perception of microbe-associated molecular patterns (MAMPs) (Hou *et al*, 2021). For example, PrePIP1 and PrePIP2 are induced by the bacterial MAMPs, flagelin 22 (flg22) and elongation factor 18 (elf18), as well as by the fungal MAMP chitin (Hou *et al*., 2014). Similarly, in Arabidopsis, ProPEP2 and ProPEP3, which are paralogues of ProPEP1, are induced upon MAMP perception and pathogen infection i.e. *Botrytis cinerea* and *Pseudomonas syringae pv. tomato* DC3000 (Huffaker *et al*., 2006). The processing of bioactive phytocytokines occurs via proteolytic cleavage of their precursors resulting into the release of small peptides of ca. 5 to 20 kDa. Phytocytokines are typically perceived by pattern recognition receptors (PRRs) (Gust *et al*., 2017, Huffaker *et al*., 2006, 2011) and activate downstream responses similar to MAMPs, although signaling pathways activated by MAMPs and phytocytokines are diverse and complex (Koenig and Moser *et al*., 2023). Therefore, the identification and mechanistic characterization of these signaling peptides is pivotal for the understanding of plant immune signaling.

The detection, quantification and identification of proteins larger than 20 kDa can be achieved through a number of methods. However, the challenge arises with small peptides such as the phytocytokines, particularly in the context of complex sample materials like plant tissues. In most cases, the method of choice for the identification of peptides is mass spectrometry, which is associated with significant experimental and technical challenges. In clinical applications, diagnostics and human sciences, a common method of choice is the Enzyme-Linked Immunosorbent Assay (ELISA). This method is established as a diagnostic standard for different clinical applications in fluid samples including blood, plasma, urine, saliva, etc. and is commonly used for the detection of antibodies, antigens, proteins and hormones such as tests for HIV or SARS-CoV-2. ELISA tests are commercialized immunological assays and are available as ready to use kits. ELISA assays are rapid and cost effective, which makes them amenable for automation and high-throughput screening (Alvarez, 2004, Posthuma-Trumpie *et al*., 2009, Martinelli *et al*., 2015). There are different types of ELISA assays, including the direct, indirect, sandwich and competitive ELISA (Alhajj and Farhana *et al*., 2023). The ELISA types differ in the antibodies, substrates and test conditions used (Fig.1)

**Figure 1.**
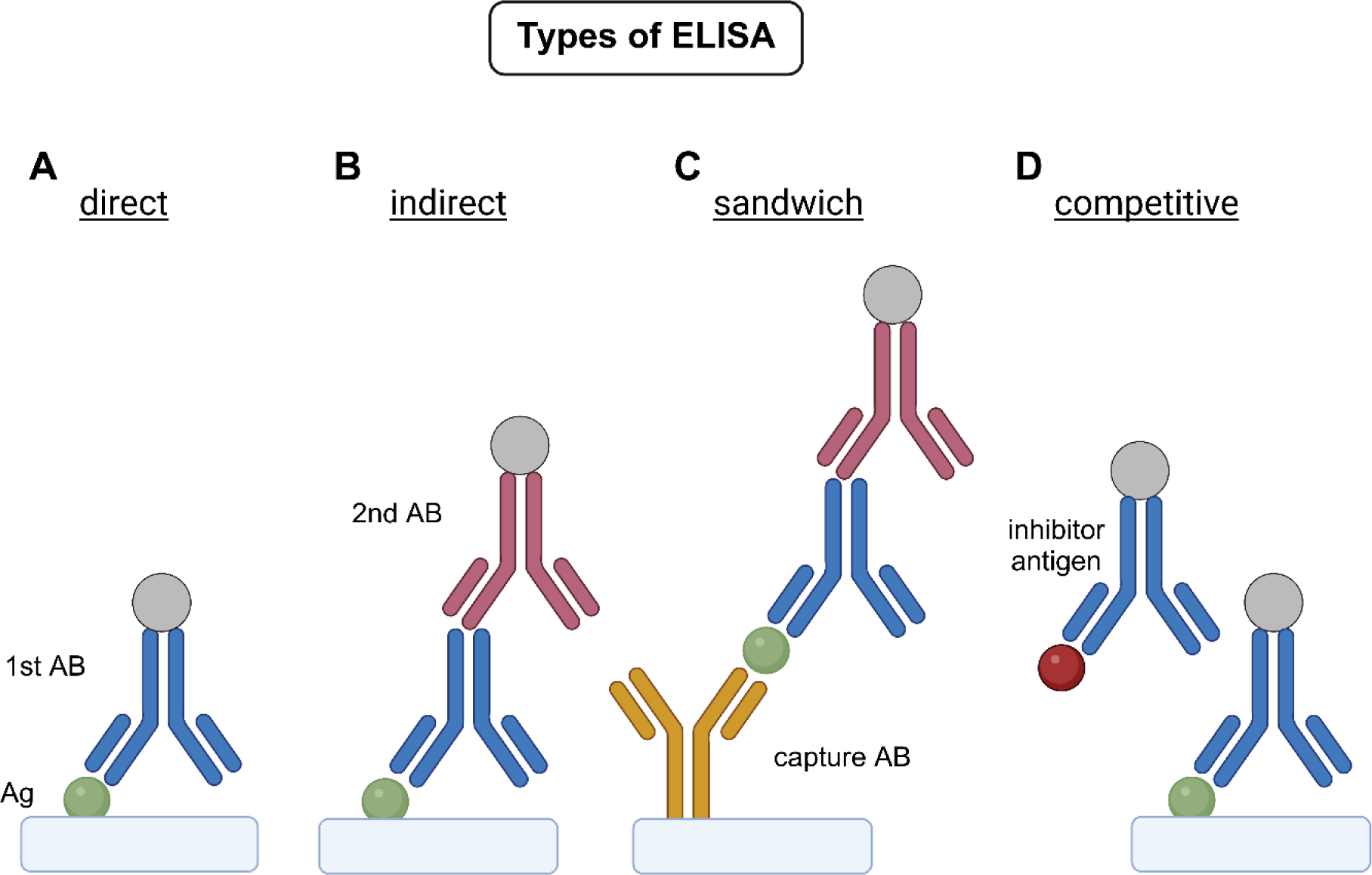
Principle and types of ELISA. Different types and basic principles of ELISA. Shown are direct (**A**), indirect (**B**), sandwich (**C**) and competitive (**D**) ELISA. Briefly, a target antigen (Ag) is bound to a surface (direct, indirect, competitive ELISA) or bound by a so-called capture antibody (AB) (sandwich ELISA). The binding of the antigen or the capture AB to a surface is referred to as “coating”. Depending on the ELISA type a primary antibody (1^st^ AB) and/or a secondary antibody (2^nd^ AB) is added and bound by the antigen (protein of interest). The detection takes place by using an enzyme linked to the first or secondary AB, for example horse radish peroxidase (HRP) [exception Immuno-PCR)]. The hydrolysis of substrate leads to the formation of products emitting a signal that is detectable by a change in relative light units (RLUs) using a luminometer, a change in optical density (OD) using a spectrophotometer or visual observation of color or a precipitated product; all indicating the presence of the antigen of interest. Graphical illustration generated with the help of BioRender®.

The antigen utilized in the ELISA is bound to a solid phase, and all ELISA types include antigen coating, blocking, detection and a final read-out (Fig.1). The direct ELISA is the simplest, fastest and less error-prone assay as there is no cross-reactivity of secondary antibodies. However, the direct ELISA is less flexible, less sensitive and less specific compared to other types, since each target requires a specific primary antibody and a secondary antibody is not applied (Fig.1A). In contrast, the indirect ELISA has the advantage over the direct ELISA in terms of sensitivity and specificity. This is due to the fact that multiple enzyme-labelled secondary antibodies bind with high specificity to the primary antibody, resulting in the amplification of the signal (Fig. 1B; Katikireddy and O’Sullivan, 2011). In an indirect ELISA, the immobilization of the antigen is not specific and therefore non-specific proteins, as in the case of a complex biological sample, may adhere to the surface of the microtiter plate (matrix) and interfere with the binding of the antigen. If the amount of antigen in a complex sample is low, indirect ELISA may not work and cross-reactivity with non-specific antigens may occur due to the use of the secondary antibody (Fig. 1B). A sandwich ELISA can be either direct or indirect, depending on the antibodies used. If the capturing antibody and the primary antibody are identical, the set-up corresponds to a modified direct ELISA. If the capturing antibody is a third and therefore different antibody, it is an indirect sandwich ELISA. A sandwich ELISA requires the immobilization of a capture antibody, which makes the binding of the antigen specific (Fig.1C). As a result, purification of the antigen from a complex biological sample may not be necessary, making the sandwich type advantageous when the antigens are present at low concentrations and/or are overlaid by non-specific antigens. However, a sandwich ELISA requires two antigen-binding sites, which might not be available, particularly in very short peptides. Furthermore, the sandwich ELISA is the most expensive and error-prone method due to the number of steps involved. It is also the slowest type of ELISA. In the competitive ELISA, also known as inhibition ELISA, the concentration of the target antigen is determined by detecting signal interference. The antigen in the sample competes with a labelled reference or standard for binding to a limited amount of antibody immobilized on the plate (Fig. 1D). Competitive ELISAs are considered for the detection of small analytes such as lipids since they allow the detection of analytes in complex mixtures including plasma, serum or cell extracts. Competitive ELISAs require less sample processing, as they are less sensitive to sample dilution and sample-related matrix effects. In addition, low inter- and intra-assay variations are observed for competitive ELISAs.

In this study, we present a method for the detection and quantification of the maize phytocytokine Zip1 in complex samples. To enable the detection of Zip1 peptides in plant material, we established an indirect ELISA-based assay (Fig. 1B). We have selected and tested experimental conditions and materials for which no commercial kits are available. The customized ELISA method can detect and quantify small peptides produced in both *in vivo* and *in vitro* samples. This method allows the highly sensitive and quantitative detection of the Zip1 peptide in leave material. By using our ELISA customized Zip1 detection we demonstrated and quantified the apoplastic release of this phytocytokine in complex plant samples under native conditions. We propose that the presented method can be broadly applied for the quantitative detection of small peptides from plant tissues.

## Material & Methods

### Generation of maize protoplasts

Grains of maize (*Zea mays, cv. Golden Bantam* or *cv. Ky21*) were placed in organic growth soil and grown for three days in a plant growth chamber (15 h photoperiod at 28°C, 1 h twilight morning and evening and 7 h darkness at 22°C). After three days (when germlings tips broke through the soil surface) the pots were placed in total darkness and grown for seven more days in the same growth chamber. 2^nd^ leaves of ten days-old etiolated seedlings were used to generate protoplasts. Leaves were piled and cut with a sterile razor blade into thin strips (approximately 1 mm). Cut material was transferred into the enzyme solution for a pre-incubation. All buffers and solutions need to be sterilized. MES was added prior use and was sterile filtrated. For protoplasting the enzyme solution was prepared containing 1% cellulose R10 (w/v) (Duchefa Biochemie), 0.2% macerozyme R10 (w/v) (Duchefa Biochemie), 0.4 M mannitol, 20 mM KCl and 20 mM MES pH 5.7. The solution was heated to 55°C for 10 min and cooled down to room temperature (RT) prior the addition of 10 mM CaCl_2_ and 0.1% bovine serum albumin (BSA) (w/v). After all material was cut it was transferred into a sterile 250 mL flask and enzyme solution was infiltrated using 60 mbar vacuum for 30 min. Subsequently, flasks were incubated for three hours at 28°C in the dark while shaking (50 rpm). Afterwards the cell suspension was filtered through a 40 µM nylon mesh (cell strainer from Falcon) into a 50 mL tube. Tubes were centrifuged for three minutes at 100 g in a swing rotor to pellet the protoplasts. The supernatant was removed and 5 mL of W5 buffer was added. W5 wash solution is made of 154 mM NaCl, 125 mM CaCl_2_, 5 mM KCl and 2 mM MES pH 5.7. Centrifugation was repeated, supernatant discarded and 5 mL W5 added. The protoplasts were incubated for 30 minutes on ice. During this incubation the protoplasts were counted using a Neubauer counting chamber. Four big squares were counted, and the amount calculated using the following equation:

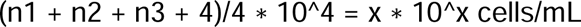

Protoplasts were pelleted (100 g, 3 min) and the pellet was resuspended in the calculated amount of MMg solution, which contains 0.4 M mannitol, 15 mM MgCl_2_ and 4 mM MES pH 5.7. 2*10^5^ cells/mL were used for the transfection.

### PEG-mediated transfection of maize protoplasts

Maize protoplasts were transfected by PEG-mediated transformation. 50 µg of column-purified plasmid (DNA template) were added to a 50 mL tube as well as 1 mL of protoplasts (2*10^5 cells/mL). Both were mixed with caution. 1.1 mL of PEG solution was added and mixed thoroughly to start the transfection process. 10 mL of PEG solution were prepared by adding 4 g of PEG4000 (Fluka, #81240 SIGMA), 3 mL of distilled water, 2.5 mL 0.8 M mannitol and 1 mL 1 M CaCl_2_. Samples were incubated for 5 minutes at RT before 4.4 mL of W5 were added to stop the transfection. Samples were pelleted (100 g, 3 min) and supernatants were carefully removed. Protoplast pellets were resuspended in 1 mL WI that contains 0.5 M mannitol, 4 mM MES pH 5.7 and 20 mM KCl. The cell suspensions were then transferred to a 12 well cell culture plate using cut 1mL tips. Samples were incubated for 16h in the dark at RT to facilitate the protein expression.

### Protoplast sample preparation

Protoplasts were harvested by using a cut tip and were transferred to 1.5 mL tubes. And pelleted by a centrifugation at 1000 g for 3 min. Afterwards the supernatant was discarded and 50 µL of prepared coating buffer was added to the pellets. 100 mM coating buffer pH 9.6 contains 0.303g Na_2_CO_3_, 0.6g NaHCO_3_ ad 100 mL distilled water. The coating buffer was supplemented with protease inhibitors supplemented with cOmplete™,Ref. 04693159001 Merck, protease inhibitor cocktail). Cell disruption was performed using an ultrasonic water bath (VWR). Samples were placed into the bath for 3 x 3 min and exposed to the highest degree of ultrasonic. Ice, was added to the water bath to keep the temperature close to 4°C. Afterwards, extracts were pelleted using a table top centrifuge at maximal speed. Supernatants were transferred to new 1.5 mL tubes and subjected to the ELISA assay.

### Preparation of *Zea mays* leaf samples

Grains of maize (*Zea mays, cv. Golden Bantam* or *cv. Ky21*) were placed in organic growth soil and grown for eight days in a plant growth chamber (15 h photoperiod at 28°C, 1h twilight morning and evening and 7 h darkness at 22°C). 2^nd^ leaves were infiltrated with distilled water (mock), 2 mM salicylic acid (SA, SIGMA Catalog No. S7401-500G) or 5 µM Zip1 peptide solution using a vacuum pump and desiccator (5 x 130 mbar, 10 min). Leaves were dried and placed in square petri dishes with wet paper tissues and incubated for 9 h. Afterwards leaf material was frozen by using liquid nitrogen and ground with a mortar and pestle. The powder was dissolved in ELISA coating buffer supplemented with protease inhibitors. For the total extract samples, 50 µL of supernatants were directly subjected to the ELISA plate for coating. For the fractionation (HMW/LMW), the supernatants were filtrated using 5 kDa molecular weight cut-off concentration columns. 50 µL of the resulting samples (flow through and supernatant) were used for coating.

### ELISA-based assay

<u>Coating</u>: Prepared sample were added to the selected ELISA-suitable antigen-binding 96-well microtiter plate (CORNING COSTAR 9018; Thermofisher scientific Ref: 44-2504-21). The first two columns contained the peptide standard, which was created using different concentrations of synthetic Zip1 (QPWEGESEKLATQGASVR) peptide (500, 250, 100, 50, 10, 5, 1, 0 ng). Samples and standards were always measured in duplicates and protein concentration was normalized either to cells/mL (protoplast) or to fresh weight (leaf samples). The loaded plate was sealed with microtiter film and for coating it was incubated overnight at 4°C. Next, coating solution was discarded and wells were three times washed with 200 µL of 1x PBS buffer pH 7.4 using a multichannel pipette. 1x PBS contains 1,44 g Na_2_HPO_4_, 0.2 g KCl, 0.246 g KH_2_PO_4_ ad 1000 mL distilled water. It is recommended to wash with pressure to remove unbound sample and buffer residues. During each wash the plate is shook for about 30 seconds and washing solution was removed by flicking the plate with force above a sink and tapping the plate onto a paper towel until the paper remained dry.

<u>Blocking</u>: 200 µL blocking solution (1x PBS, 5% skim milk or BSA) was applied subsequently to block-remaining protein-binding sites. The plate was again sealed and incubated for 3 h at RT. After blocking, it is optional to wash the plate as described after coating. Additional washing reduces background, but may decrease the sensitivity.

<u>Detection</u>: 100 µL of 7 µg/mL custom α-Zip1 antibody was added to the wells. The antibody was prepared in 1xPBS 5% skim milk and it is recommended to dilute the antibody fresh prior use, since reusing may affect the assays sensitivity. The plate was sealed and incubated overnight at 4°C. Afterwards the plate is washed four times as previously described. Next, 100 µL of the secondary antibody (α-rabbit-HRP 1:1000; Cell Signaling #7074) was added to the wells and the plate was sealed and incubated for 2 h at RT. For the detection, the plate was washed four times as described before and 100 µL of cover solution (4% bromphenolblue [w/v] dissolved in 70% EtOH) was added carefully to empty wells. Subsequently, 50 µL bubble-free Femto solution (SuperSignal^TM^ ELISA Femto Maximum Sensitivity Substrate; Ref. 37075) was added to the wells. The plate was placed into a luminometer (TECAN Infinite M200Pro, Software: Tecan i-control v.1.11.1.0) and the chemiluminescence was measured. A standard curve was generated for each plate from the serial dilution (value of blank is deducted). Antigen concentrations of samples were extrapolated from the standard curve.

## Results

In this study, we have established an indirect ELISA for the detection of the maize phytocytokine Zip1 (Fig.1B). Therefore, a custom α-Zip1 antibody was synthesized and tested regarding its specificity, selectivity and sensitivity. First, different concentrations of recombinant PROZIP1 (rPROZIP1) were monitored using 1 µg/mL α-Zip1 antibody. Up to 12 ng of rPROZIP1 signals at 17 kDa corresponding to full length PROZIP1 were observed with the α-Zip1 antibody compared to PROZIP1 visualized by SYPRO® Ruby, which showed signals up to 25 ng, indicating that the α-Zip1 antibody is sensitive to Zip1 (Fig. 2A). To test for the specificity and selectivity of the α-Zip1 antibody, mcherry-PROZIP1-GFP and a Zip1-containing cleavage site mutant CS1 (PROZIP1_CS) (Ziemann *et al*., 2018) were heterologous expressed in *N. benthamiana*. GFP served as a control. Different volumes of total protein extract were tested and detected using the α-Zip1 antibody. Signals specific for PROZIP1, PROZIP1_CS and putative Zip1-containing proteins were observed (Fig. 2B). No signals in GFP were detected indicating that the antibody is specific for Zip1. Next, PROZIP1-GFP and PROZIP1_CS-GFP were expressed in maize protoplast of the cultivar *Golden Bantam* (*GB*). GFP served as control and was expressed in the cultivars *GB* and *Ky21*, of which the latter does not contain PROZIP1. Using the α-Zip1 antibody for detection, signals in PROZIP1-GFP and PROZIP1-CS-GFP samples of *GB* were detected, but not in GFP samples of *GB* and *Ky21* confirming the specificity of the α-Zip1 antibody (Fig. 2C).

**Figure 2.**
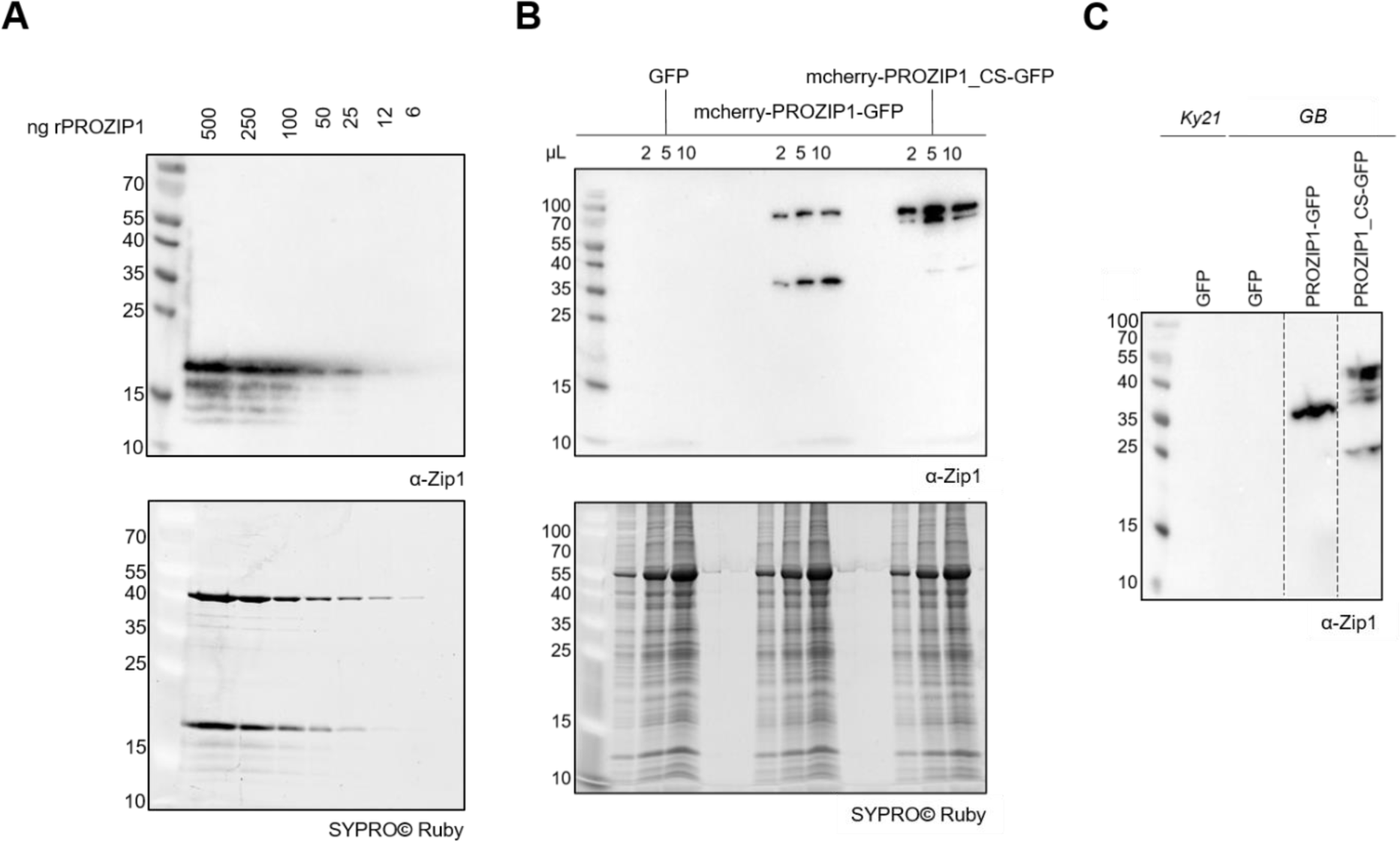
Zip1 antibody test. Zip1 antibody was tested for specificity, selectivity and sensitivity. (**A)** Different amounts of recombinant PROZIP1 (rPROZIP1) were detected by α-Zip1 antibody (1 µg/mL). As a control SYPRO® Ruby staining was performed (**B)** Overexpression of GFP, mcherry-PROZIP1-GFP and mcherry-PROZIP1_CS-GFP in *N. benthamiana* using Agrobacterium-mediated transformation. 2, 5 or 10 µL of total protein extract (100 mg) was tested by SYPRO® Ruby staining and western blot using the α-Zip1 antibody (7 µg/mL). **(C)** GFP, PROZIP1-GFP and PROZIP1-CS-GFP was overexpressed in *Z. mays* protoplast using PEG-mediated transformation. GFP was expressed in both maize cultivars *Ky21* and *Golden Bantam* (GB) and detected by western blot using the α-Zip1 antibody (7 µg/mL).

For the adapted ELISA assay, synthetic Zip1 antigen and antigen from complex biological samples are coated onto the plate and bound by a specific α-Zip1 antibody. α-Zip1 antibody signals are enhanced by binding to an α-rabbit-HRP secondary antibody. The detection utilizes ECL-substrate, which is oxidized by HRP to generate a luminescent signal (Fig. 3).

**Figure 3.**
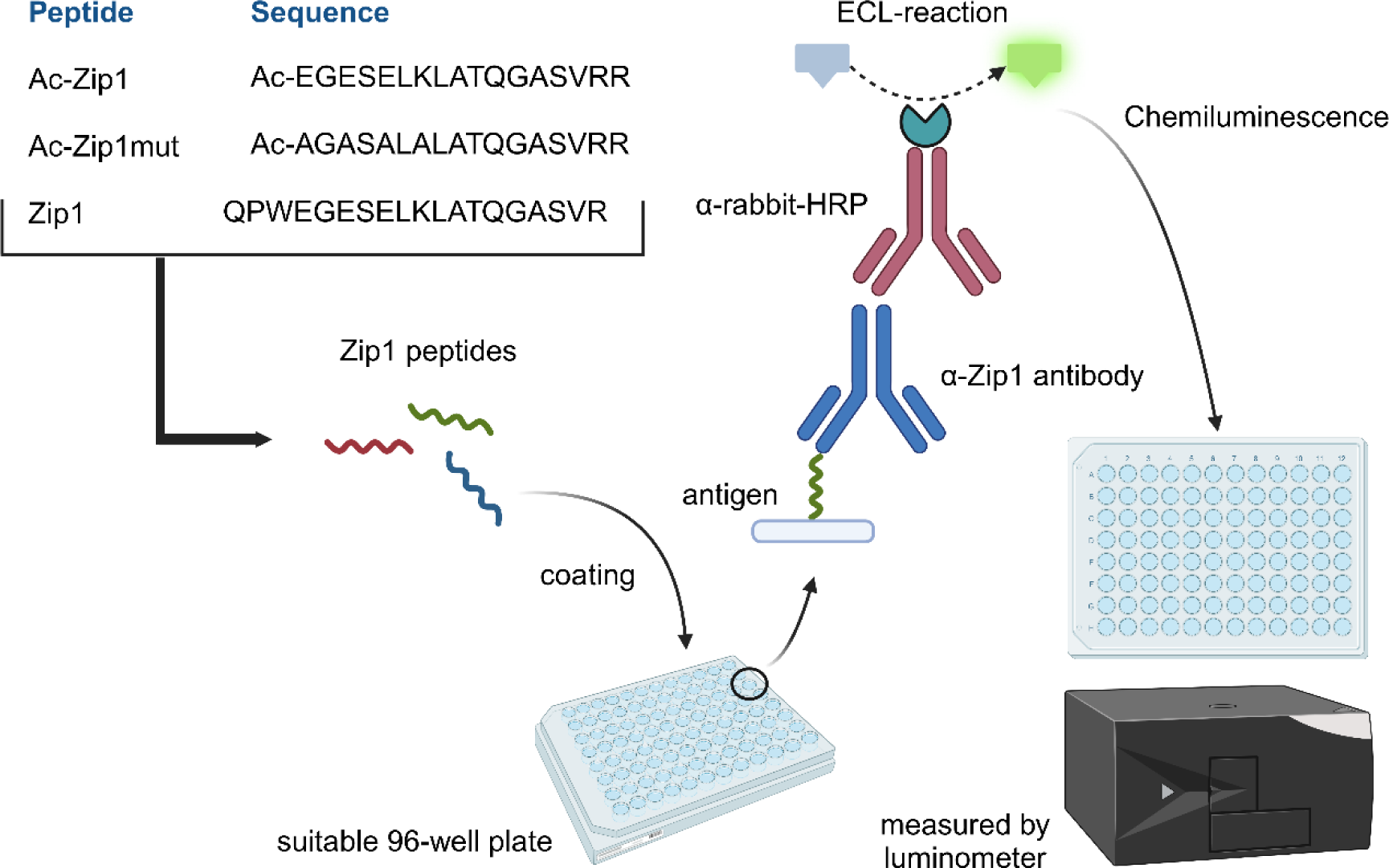
Schematic overview of the adapted ELISA assay for the detection of Zip1 peptide. Synthetic Zip1 peptides were used as antigens and coated to the ELISA plate for standardization. The coated plate was subjected to an established α-Zip1 antibody. The first antibody is bound by a secondary HRP-conjugated α-rabbit antibody (indirect ELISA). HRP-mediated ECL (enhanced chemiluminescence) reaction leads to emerging chemiluminescence, which is measured by a luminometer.

To establish the ELISA-based assay, different measurements were carried out in order to adjust the selectivity, sensitivity, specificity and reproducibility of the assay. For the sensitivity, a range of α-Zip1 antibody concentrations was tested, with the lowest concentration of 2.5 µg/ml α-Zip1 antibody proving to be the least amount needed to measure a signal (Fig. 4A). A concentration of 7 µg/ml of the α-Zip1 antibody was selected for further experiments, as this provides the most effective signal intensity within the linear signal/concentration range (Fig. 4A). To test for sequence specificity, the synthetic variants Zip1 (EGESELKLATQGASVR) and Zip1-mut (AGASALAKLATQGASVRR) were tested. Luminescence signals were detected in a concentration-dependent manner for Zip1, but not for the Zip1-mut peptide (Fig. 4B), demonstrating binding specificity of the α-Zip1-antibody and selectivity against the Zip1 epitope. The quantification of the signals resulted in a detection limit of 5 ng/mL of Zip1 antigen in contrast to the Zip1-mut, which was only detectable at concentrations above ≥500 ng/mL peptide, indicating the high specificity of these ELISA assay for the Zip1 antigen (Fig. 4C). Besides, different amounts of bovine serum albumin (BSA) were tested to mimic background proteins in complex samples. The results did not show a significant increase in signal compared to Zip1 antigen, suggesting that the tested concentrations of non-Zip1 protein did not interfere with the specificity and selectivity of the α-Zip1 antibody (Fig. 4D). For the standardization of the assay, a calibration curve was generated using a concentration range from 1 to 1000 ng/mL Zip1 peptide. The curve aligns along the measurements resulting in a coefficient of determination (R^2^) value of 0.9904 (Fig. 4E). This standard curve was generated for each ELISA assay. In addition to luminescence detection, we attempted colorimetric detection and tested the 3,3’,5,5’-tetramethylbenzidine (TMB) reagent (Ref. T444 Sigma-Aldrich) to compare the luminescence and the colorimetric detection. However, due to the extremely low coloration observed in the tested samples, the colorimetric detection did not yield sufficient data points to generate an effective linear standard curve (R^2^=0.5416) (Fig. 4F).

**Figure 4.**
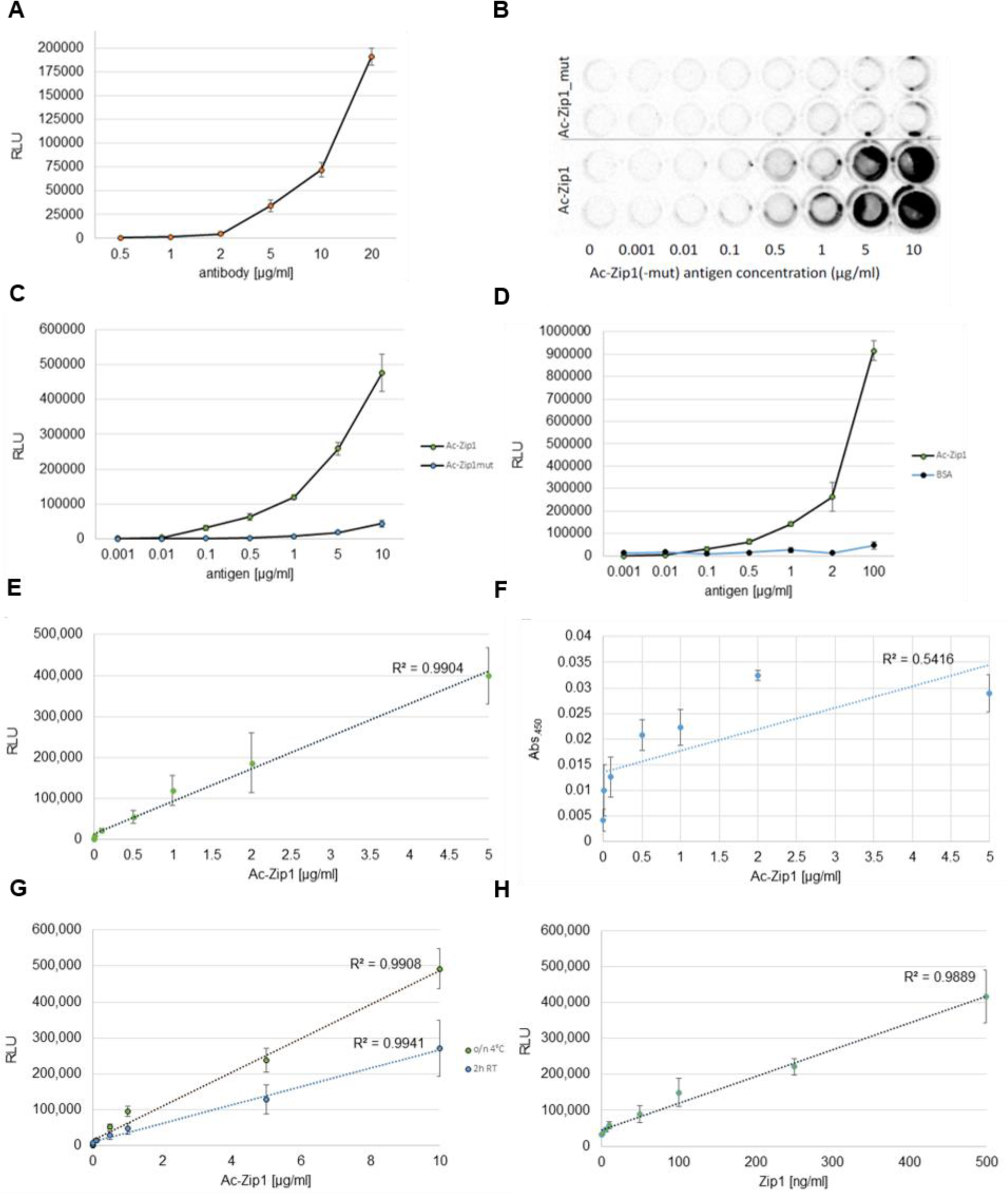
Optimization of the most effective conditions for the Zip1 detection. **(A)** Optimal α-Zip1-antibody concentration **(B-C)** α-Zip1-antibody specificity and selectivity using synthetic N-terminally acetylated (Ac) Ac-Zip1 peptide and Ac-Zip1mut **(D)** Antibody specificity using the example of bovine serum albumin (BSA) **(E)** Standardization using ECL substrate **(F)** Standardization using colorimetric TMB reagent **(G)** Coating overnight in cold vs. 2h room temperature (RT) coating **(H)** Final standard using Zip1 as antigen while coating overnight in cold and ECL driven-detection. RLU= Relative Light Unit, Abs=Absorption.

For the optimization of the coating, we tested two different conditions, coating overnight at 4°C and coating for two hours at RT. Considering the R^2^ values of the overnight coating (0.9908) and the two-hour coating (0.9941), both conditions lead to similarly good results. For further experiments we used overnight coating as the measured RLU range was higher compared to the two-hour coating, suggesting that more antigen was bound to the microtiter plate, which might increase the sensitivity (Fig. 4G). Two different Zip1 peptides were tested as antigens including the N-terminally acetylated (Ac-) Ac-EGESELKLATQGASVRR and the non-acetylated QPWEGESELKLATQGASVR (Fig. 1 & Fig. 4H). The 17 amino acid Zip1 peptide EGESELKLATQGASVRR was identified in SA-treated maize leaves (Ziemann *et al*., 2018) and the synthesized peptide was N-terminal acetylated. Peptide modifications such as acetylation of the N-termini can increase peptide stability and are a closer mimic of native conditions (https://www.lifetein.com/Peptide-Synthesis-Amidation-Acetylation.html; Arispe *et al*., 2008). Furthermore, an acetylation might increase the biological activity of a peptide and reduce the effect of charged C- or N-termini during ELISA binding (Arispe *et al*., 2008). In addition, the 19 amino acid peptide QPWEGESELKLATQGASVR was included as it may result from the proposed arginine-dependent processing of PROZIP1 (Ziemann et al., 2018). Comparing the tested Zip1 antigens, the peptide QPWEGESELKLATQGASVR delivered the best standard curve concerning variability and a better resolution at the low nanogram range (Fig. 4H), suggesting that the increased peptide length and amino acid variation lead to an increased peptide adhesion.

As a proof of concept, we tested the sensitivity of our method to monitor Zip1 in biological samples. To this end we transfected maize protoplasts with constructs overexpressing the precursor of Zip1, PROZIP1 containing a C-terminal GFP-tag. We only observed signals for PROZIP1-GFP samples compared to the GFP control samples in transfected protoplasts (Fig. 5A). Signals in the GFP background control were significantly low reflecting the specificity of the ELISA towards Zip1. Consequently, all values were normalized to the GFP signals control. Since Zip1 has been first detected in the apoplast of SA-treated maize plants (Ziemann *et al*., 2018), we used SA as trigger to induce the natural Zip1 production in maize *cv. GB* leaves. In addition, leaves were treated with Zip1 peptide, since it is known from several phytocytokines that the presence of the stress signal might lead to an induction of expression of the corresponding precursors and likely release of phytocytokines (Huffaker *et al*. 2011). Maize *cv. Ky21* served as a background control, which does not contain the *prozip1* gene. Hence, *Ky21* mock signal values were used for normalization. SA and Zip1-treated samples showed a significant signal increase compared to *GB* mock samples and *Ky21* samples (Fig. 5B). Interestingly, Zip1 treatment resulted in higher signal detection compared to SA, which could be caused by the self-detection of the Zip1-treated peptide. *Ky21* did not show a significant signal increase within treatments (Fig. 5B). Signal quantification showed approximately 60 ng/mL of PROZIP1 or Zip1-containing peptides detected in SA-treated samples compared to approximately 100 ng/mL detected by the Zip1 treatment (Fig. 5C), suggesting a higher production of Zip1-containing peptides in the Zip1-treated sample or a self-detection of the treated Zip1 peptide. These results demonstrate a natural detection of Zip1-containing fragments using our ELISA method.

**Figure 5.**
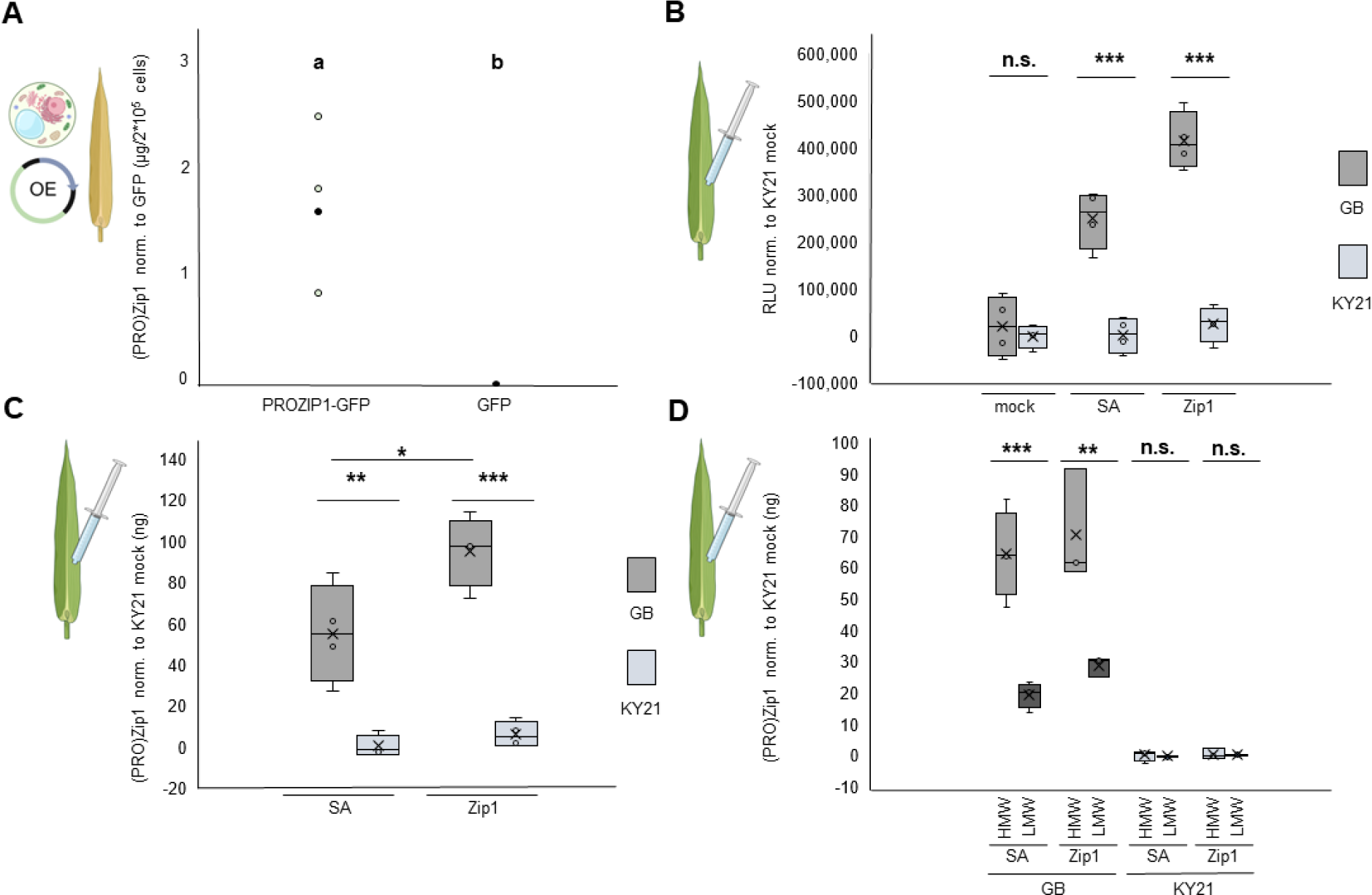
Increased abundance of Zip1-containing peptides after SA and Zip1 treatments in *Z. mays*. **(A)** Protoplasts of *cv. Ky21* were transfected using an overexpression (OE) construct of PROZIP1-GFP. GFP signals served as a control. Leaves of two maize cultivars (*cv. GB* and *Ky21*) were treated with 2 mM SA or 5 µM Zip1 peptide and incubated for 9h. Proteins were extracted in coating buffer (CB) supplemented with protease inhibitors from ground leaf powder and subjected to the ELISA-based assay. **(C)** Quantification of Zip1-containing peptides. Values were normalised to the *Ky21* mock signals. **(D)** Total extract (TE) of SA and Zip1 treated samples was fractionated by molecular weight (low molecular weight (LMW = <5 kDa, high molecular weight (HMW = >5 kDa). The experiments were performed with at least three independent biological replicates. Statistical analysis was performed using an ANOVA or Students t-test. Letters or asterisk above indicate significant differences between samples (p=0.05*, 0.01*, 0.001*).

To differentiate between signals from full-length PROZIP1 (15 kDa) and Zip1 peptide (1 kDa), supernatants were separated using concentration columns with a molecular weight cut-off of 5 kDa and monitored via the ELISA assay. The high-molecular weight fractions (HMWF, >5 kDa) showed signals of around 60 ng/mL whereas the low-molecular weight fractions (LMWF, <5 kDA) resulted in signals of around 25 ng/mL, suggesting a higher detection of longer fragments containing Zip1 than the bioactive Zip1 peptide itself (Fig. 5D). No signals were detected in fractions of *Ky21* reflecting the specificity of the ELISA and indicating that the Zip1 treatment does not lead to the detection of Zip1 peptide itself (Fig. 5D). Overall, our results indicate that SA and Zip1 treatments resulted in an increased level of Zip1-containing peptides, particularly in the Zip1-treated sample. *In vivo* we were able to detect approximately 25 ng/mL of Zip1-containing peptides in the low molecular weight fractions indicating this amount of Zip1 present in our tested biological samples. Considering the plant material used and the enrichment of small peptides, the measured quantity represents the actual minimal amount of bioactive Zip1: 1.66 µg of Zip1 is present in 1 g of SA- or Zip1-treated leaf material, which corresponds to approximately 80 nM Zip1. Finally, we show that the Zip1-ELISA is a specific and reliable method to monitor the abundance of Zip1-containing peptides *in vivo*.

## Discussion

In this study, we present a customized ELISA protocol that allows sequence-specific and quantitative detection of small peptides from native maize tissue. In general, ELISAs (commercial and custom) are commonly used in agricultural and nutritional sciences to detect allergens and plant pathogens. A partially custom-made ELISA assay by Liu *et al*. (2021), targeting the detection of the soybean alpha-subunit of beta-conglycinin reached a detection limit of 0.65 ng/mL. Monaghan *et al*. (2008) utilized a custom-made indirect ELISA to detect sulphur-rich protein (SRP) in soybean meeting our detection limit of 5 ng/mL. Other than that, ELISAs are used to determine the content of prolamin (gluten precursor) in foodstuffs (Diaz-Amigo *et al*., 2013). It should be noted that the use of ELISAs for the quantification of prolamin (gluten) is under debate and other methods such as LC-MS are being discussed. However, ELISA is still the method of choice for prolamin quantification in foodstuffs (Alves, T. O. *et al*., 2019). For the detection of plant pathogens, ELISA assays are recommended by the European and Mediterranean Plant Protection Organization (EPPO) to test for the presence of viruses on grafting material of fruit trees (Boonham *et al*., 2014). In addition, new ELISA assays are developed for the detection of plant pathogens. Gorris *et al*. (2020) described the development of a so-called double antibody sandwich ELISA (DAS-ELISA) for the detection of the bacterial pathogen *Xylella fastidiosa*. DAS-ELISA assays appeared to be very specific, as no false-positive results were observed in field samples. In addition, DAS-ELISA assays are reliable with a sensitivity of 104 colony-forming units (CFU) per mL. Wu *et al*. (2011) described the development of a triple antibody sandwich ELISA assay using a monoclonal antibody specific for the detection of Odontoglossum ringspot virus, which successful demonstrated no cross-reactivity in samples challenged with seven other viruses.

There are several factors to consider when setting up an ELISA assay. The sensitivity and specificity of ELISA can vary considerably between different assays. This variability is largely dependent on the ELISA type, the antibodies used (monoclonal or polyclonal), the conjugated enzyme and the corresponding substrate (Venbrux *et al*., 2023). In addition, the choice of the correct plate (matrix) has a major impact on the specificity and sensitivity of ELISA, a problem commonly referred to as “matrix interference” (Wang *et al*., 2007). Depending on the type of antigen, i.e. lipid, protein, glycan, there is a wide range of matrices available. The surface of such matrices is usually biochemically modified to enhance the potential binding of the desired antigen. Knowledge of the biochemical properties of the antigen, especially its hydrophobicity/hydrophilicity is therefore crucial. In the case of Zip1 we chose a matrix, which is designed to bind a wide range of biomolecules with a tendency to bind preferentially slightly hydrophobic compounds. We used a custom α-Zip1 antibody designed against the complete Zip1 peptide sequence of the mature phytocytokine (EGESELKLATQGASVRR, Ziemann *et al*., 2018). Because the α-Zip1 antibody was tested regarding its specificity, selectivity and sensitivity, we chose an indirect ELISA to profit from the higher sensitivity and selectivity, while taking the risk of low detection knowing the naturally low abundance of Zip1 antigen in complex biological samples.

In addition to the surface, the desired detection type contributes to the selection of a suitable matrix. Most ELISA assays use colorimetric or luminescent readouts. As the name suggests a colorimetric detection produces a color as a readout and therefore a transparent or non-transparent matrix (plate) is used for the detection. In contrast, a non-transparent plate should be used for luminescent detection to avoid signal scattering. In our hands, the transparent ELISA plate worked best with the Zip1 antigen coating and we had to overcome signal scatter with an opaque cover solution. In addition, we loaded samples onto the plate at the greatest possible distance from other samples to simulate an opaque plate. We were unable to use TMB as a colorimetric substrate in our assay as the detection limit was not achieved in our standard curves, resulting in inadequate R^2^ values. Similar to ECL substrate, the TMB reaction is catalyzed by HRP in a redox reaction and in case of initially colorless TMB this results in a slight blue product that has an absorption of 370 to 655 nm. The reaction is stopped by adding 2 M H_2_SO_4_ and results in a yellow color and an absorption at 450 nm. Because no significant color was observed, we suspected a low reactivity between the used TMB substrate and the HRP as our antibody is custom-made. The successful establishment of our ELISA-based assay is reflected by the R^2^ values obtained for the standard curves as R^2^ values >0.9 are good indicators of signal quantification (Quian *et al*., 2009, Rampitsch *et al*., 2003). The use of protease inhibitors during sample preparation and coating is highly recommended to avoid and minimize variations in the peptide quantification (Aydin *et al*., 2015). In general, if possible, we can recommend purification steps of the antigen of interest to improve the specific signals while reducing the background. On the other hand, immunoassays such as ELISA are recognized as the most powerful method for analysis of large amounts of plant material due to their accuracy and simplicity (Yoshimatsu *et al*., 1995). If the affinity of an antibody for its antigen is sufficiently high, the antibody can also bind its antigen in crude extracts (Yoshimatsu *et al*., 1995). While there are numerous benefits to using an ELISA-based method in plant tissues, there are also drawbacks to be considered. For example, due to the chemical and physical stability of antibodies, they require refrigeration for storage and, in some cases, special buffers. In addition, the production of novel antibodies can be complicated and expensive (Sakamoto *et al*., 2018).

Phytocytokines are signaling molecules with an overall low abundance. The exact cellular concentration is not currently known for any phytocytokine (Rhodes *et al*., 2021). Therefore, detecting phytocytokines using an ELISA-based assay is not a guarantee for success. The ratio of antigen (phytocytokine) to non-specific background is not favorable. In addition, a high specific and selective primary antibody is mostly essential. We do not recommend performing our assay using tagged proteins as the risk of fishing non-specific signals is much higher. Nevertheless, this assay can be an excellent choice compared to other limited, expensive and time-consuming alternatives such as mass spectrometry, which is the gold standard for the identification of peptides from complex samples. In addition to the high demands in equipment, consumables and labour, mass spectrometry also relies on the quality of available databases, which can be a critical bottleneck when hunting for specific small peptides. Similar to ELISA, we attempted to identify Zip1 and Zip1-containing peptides in the low molecular weight samples by mass spectrometry, but were unsuccessful. Using mass spectrometry there is still the possibility to not identify the searched peptide as it can be modified e.g. post-translational modifications (PTMs), it cannot “fly” e.g. fail to ionize or the ions are too unstable and fragmentate, it is conserved between different proteins or it is not present in the database of interest. In contrast, an ELISA-based assay can be advantageous since the actual peptide is detected independent of its modification by a very specific biochemical reaction (antigen-antibody binding).

Using the ELISA-based assay, we were able to detect native Zip1 peptide in *Z. mays* leaf samples. Salicylic acid and the presence of synthetic Zip1 triggered the native expression of prozip1 *in vivo* thus allowing the detection of an increase of Zip1-containing peptides. The application of both triggers, SA and Zip1, might initiate the processing and release of native Zip1 *in vivo*, as more fragments were quantified in the low molecular weight fractions. Since several protease inhibitors were used during and after the extraction processing and release occurred *in vivo*.

We have designed and optimized ELISA to enable the detection and quantification of phytocytokines in plant material. Using the adapted method, we were able to detect approximately 80 nM of Zip1 in SA- and Zip1-treated maize leaves. In future, the presented method may help to detect other known phytocytokines and to provide insight into their processing and release mechanisms as our method is versatile and can be used in diverse plant systems.

## Acknowledgments

We acknowledge support from the Deutsche Forschungsgemeinschaft (DFG, German Research Foundation) through the project DO 1421/5-2, the SFB1403 project number 414786233 as well as from the Cluster of Excellence on Plant Sciences (CEPLAS) funded under Germany’s Excellence Strategy – EXC 2048/1 – project ID: 390686111.

## Author contributions

MK, GD and JCMV designed the study. ZS and MK performed the experiments. MK wrote the manuscript with contributions from all authors.

## Conflict of interest

The authors declare no conflict of interest.

## Supplementary figures

**Figure S1.**
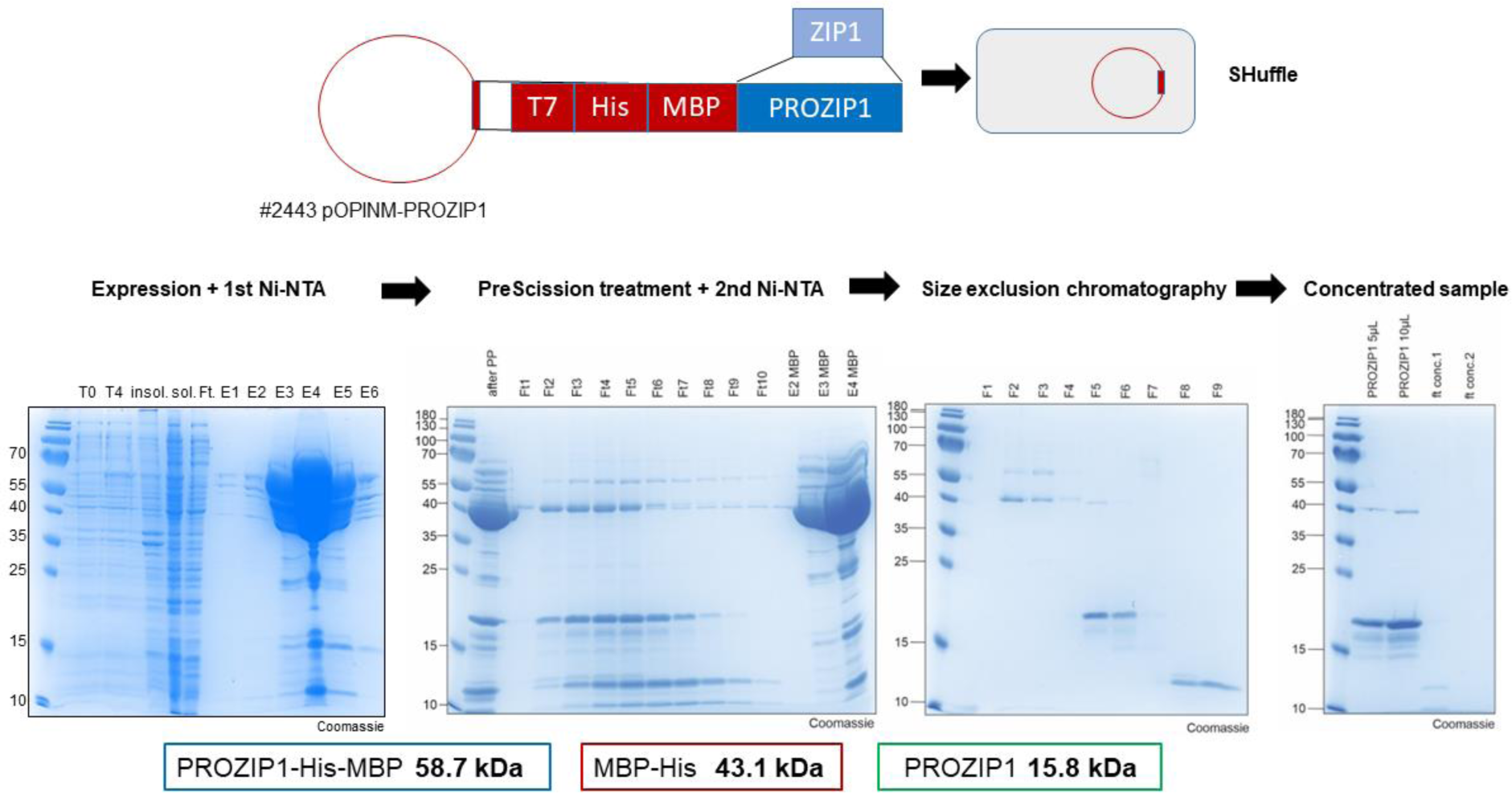
Expression and purification of recombinant PROZIP1 (rPROZIP1). SHuffle cells were used for 4 h expression at 30°C. Two Ni-NTA were performed one after another. The 1^st^ Ni-NTA was used to affinity purify PROZIP1-His-MBP. Afterwards a PreScission Protease treatment was performed to cleave between the His tag and PROZIP1 and sample was subjected to a second Ni-NTA. His-MPB was removed by the 2^nd^ Ni-NTA and the flow through collected to recover PROZIP1. To separate traces of free His-MBP from PROZIP1, the flow through was subjected to size exclusion chromatography. Finally, PROZIP1 containing fractions were concentrated using a 5 kDa molecular weight concentrator columns. The flow through and the concentrated samples were tested.

## Notes

### Competing Interest Statement

The authors have declared no competing interest.

